# Predicting protein domain temperature adaptation across the prokaryote-eukaryote divide

**DOI:** 10.1101/2021.07.13.452245

**Authors:** Sarah E. Jensen, Lynn C. Johnson, Terry Casstevens, Edward S. Buckler

**Affiliations:** School of Integrative Plant Sciences, Plant Breeding and Genetics Division, Cornell University, Ithaca, NY, 14853, USA; Institute for Genomic Diversity, Cornell University, Ithaca, NY, 14853, USA; United States Department of Agriculture, Agricultural Research Service, Ithaca, NY, 14850, USA

## Abstract

Protein thermostability is important for fitness but difficult to measure across the proteome. Fortunately, protein thermostability is correlated with prokaryote optimal growth temperatures (OGTs), which can be predicted from genome features. Models that can predict temperature sensitivity across the prokaryote-eukaryote divide would help inform how eukaryotes adapt to elevated temperatures, such as those predicted by climate change models. In this study we test whether prediction models can cross the prokaryote-eukaryote divide to predict protein stability in both prokaryotes and eukaryotes. We compare models built using a) the whole proteome, b) Pfam domains, and c) individual amino acid residues. Proteome-wide models accurately predict prokaryote optimal growth temperatures (r^2^ up to 0.93), while site-specific models demonstrate that nearly half of the proteome is associated with optimal growth temperature in both Archaea and Bacteria. Comparisons with the small number of eukaryotes with temperature sensitivity data suggest that site-specific models are the most transferable across the prokaryote-eukaryote divide. Using the site-specific models, we evaluated temperature sensitivity for 323,850 amino acid residues in 2,088 Pfam domain clusters in Archaea and Bacteria species separately. 59.0% of tested residues are significantly associated with OGT in Archaea and 75.2% of tested residues are significantly associated with OGT in Bacteria species at a 5% false discovery rate. These models make it possible to identify which Pfam domains and amino acid residues are involved in temperature adaptation and facilitate future research questions about how species will fare in the face of increasing environmental temperatures.

## Introduction

Understanding what makes a species resistant to heat stress starts with the proteome because high temperatures affect protein biochemistry, folding, and function (Fields et al. 2015; Laye et al. 2017; Ritchie 2018; Hait et al. 2020). Extreme temperatures can reduce enzyme stability and cause protein denaturation and aggregation in the cell. Protein denaturation is a problem because aggregated proteins can become cytotoxic, and because protein metabolism is energetically expensive. If proteins are too unstable, even minor increases in temperature can cause protein denaturation and force the cell to devote extra energy to protein synthesis and recycling. As a result, stable proteins confer a substantial fitness benefit to their host (Geiler-Samerotte et al. 2011; Goff 2011; Lynch and Marinov 2015). By comparing protein thermostability across species, we can gain insights into how species adapt to high temperatures and facilitate questions about how climate change might affect evolution at the molecular level.

Protein stabilities in the cell follow a distribution; the average protein is relatively stable, but there is a tail of unstable proteins that are prone to denaturation. In fact, as much as 15% of the proteome may be composed of proteins that are only stable across a temperature range of less than 5 degrees Celsius (Ghosh and Dill 2010). These marginally-stable proteins likely exist because of constraints on protein evolution. Evolution favors protein conformations that avoid complete unfolding, but purifying selection is too weak in most proteins to avoid conformations that are only marginally-stable (Razban et al. 2021). The strength of purifying selection is related to protein expression. Because protein metabolism is energetically expensive, highly-expressed proteins tend to be more stable than proteins with low or moderate levels of expression (Leuenberger et al. 2017).

Extensive studies into each protein would make it possible to predict which amino acid residues are most important for stability and predict how changing one or more residues would affect protein stability. Experimentally determining amino acid substitution effects in such a comprehensive way is, however, still too time- and resource-intensive to apply across an entire proteome. Fortunately, evolution has already run billions of independent experiments to determine which amino acid substitutions increase protein thermotolerance and organism fitness. Over time, minor residue changes compound to alter intramolecular interaction networks and adapt protein temperature sensitivity in response to the environment (Gu and Hilser 2009; Heizer et al. 2011). Comparing residue changes across species adapted to different temperatures can provide insights into how the observed amino acid mutations affect protein stability (Holland et al. 1997; Fields et al. 2006; Lockwood and Somero 2012).

Prokaryotic species have short generation times and have adapted to a wider range of optimal temperatures than eukaryotes, which makes them good species in which to study protein thermostability. They also have effective population sizes on the order of 10^8^-10^9^, which means selection can act more efficiently in these species than in most eukaryotes (Clarke 2014; Bobay and Ochman 2018). As an added benefit, there are a large number of prokaryote species with available genome and proteome sequences that can be used to develop models of protein temperature sensitivity. Prokaryote optimal growth temperatures (OGTs) can be predicted using genome features, and OGT is known to correlate with proteome temperature sensitivity (Saelensminde et al. 2007; Zeldovich et al. 2007; Dehouck et al. 2008; Gu and Hilser 2009; Jensen et al. 2012; Meruelo et al. 2012; Aptekmann and Nadra 2018; Li et al. 2019; Sauer and Wang 2019; Cimen et al. 2020). OGT predictions from tRNA sequences provide protein-independent estimates of temperature sensitivity that can then be used to evaluate protein thermal stability (Cimen et al. 2020). Models built with prokaryotic protein data that can be applied to predict eukaryotic protein thermal stabilities would help explain eukaryotic molecular adaptation to high temperatures. Models that can identify temperature-sensitive amino acid residues would make it possible to ask questions about how and why individual amino acids affect protein thermostability.

Evolutionary constraints on protein metabolism are consistent across phylogenetic domains, as is biochemistry (Camps et al. 2007; Harms and Thornton 2013; Venev and Zeldovich 2018). We hypothesize that this shared biochemistry will make it possible to also share information about protein stability across species separated by large evolutionary time scales. Residue chemistry and protein stability estimates may be consistent and predictable across the prokaryote-eukaryote divide. With this in mind, we set out to understand whether protein sequence features can be used to compare protein temperature sensitivity across all domains of life and provide insights into eukaryotic protein adaptation. Because protein structures are only available for a subset of proteins, we opted to restrict our model inputs to features of primary sequence. We developed models that test the transferability of a) proteome-wide sequence features, b) Pfam domain sequence features, or c) features of single amino acid residues. Our models are built and tested on a dataset of ∼13.8 million proteins from 4,832 species across all three domains of life spanning more than 3 billion years of evolution (Battistuzzi et al. 2004; Hug et al. 2016).

## Results

### Proteome amino acid composition predicts OGT in prokaryotes, but not eukaryotes

4,827 Archaea and Bacteria species were evaluated to see if there is a relationship between proteome amino acid content and optimal growth temperature (OGT). Consistent with previous studies, we find that prokaryotic OGT can be predicted by amino acid composition in the proteome (Zeldovich et al. 2007; Dehouck et al. 2008; Jensen et al. 2012; Sauer and Wang 2019). We find that the accuracy of OGT predictions depends on which species are used to build the model and that Pfam domain regions of the protein are the most predictive. When the model is built using Archaea species, accuracy reaches r^2^ values up to 0.37 but overestimates Bacteria OGTs (Figure 1A, Table 1). The coefficient of determination between true OGT and predicted OGT is highest when the model is built with Bacteria species (r^2^ up to 0.93), but this model tends to underestimate Archaea OGTs (Figure 1B, Table 1). The most predictive models (with both high r^2^ and low RMSE) come from using a random sample of both Archaea and Bacteria amino acid frequencies. The maximum r^2^ from these models is 0.82, and the root mean square error (RMSE) is only 4.6°C (Figure 1C, Table 1).

**Figure 1:**
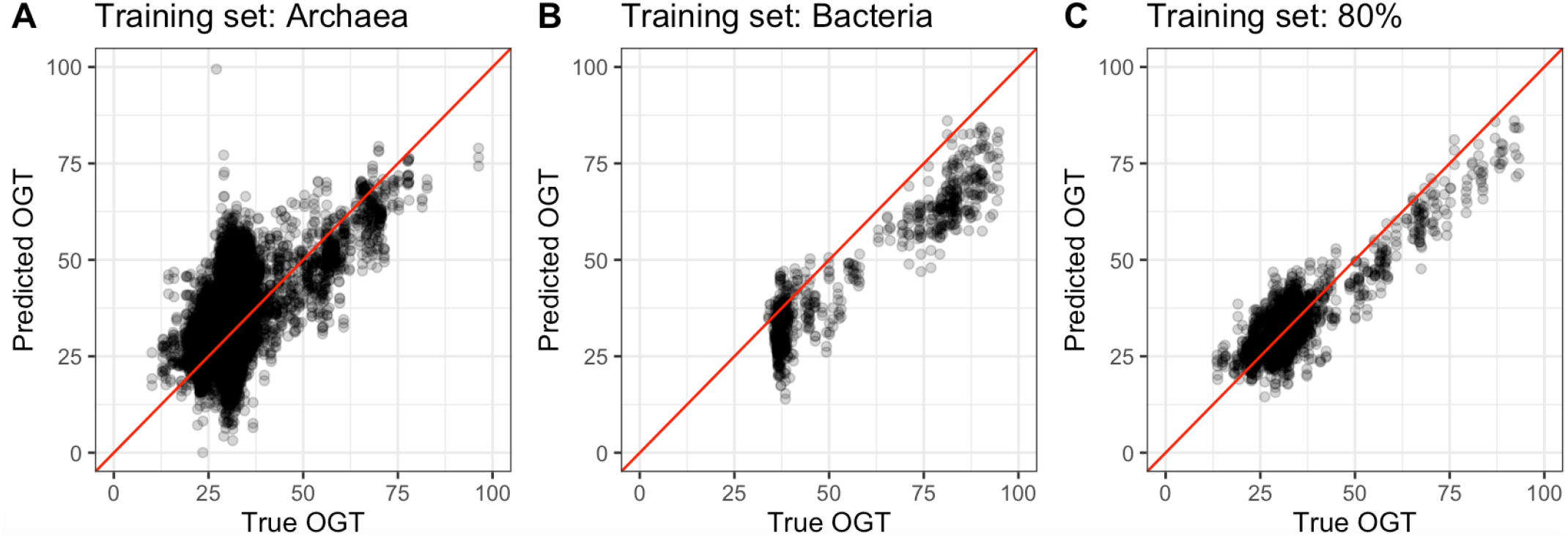
Performance of global models using amino acid proportions across the whole proteome. A) Model built with Archaea species and applied to Bacteria species. B) Model built with Bacteria species and applied to Archaea species. C) Model built with 80% of all data and applied to a held-out 20%. Red lines show the point where the true OGT values match predicted OGT values.

**Table 1:** Global model performance when trained with Archaea, Bacteria, or a combined dataset and tested on the remaining set of prokaryote species.

|  | Dataset |  |  |  |  |  |  |  |  |
| --- | --- | --- | --- | --- | --- | --- | --- | --- | --- |
|  | Whole Proteome |  |  | Pfam Domains |  |  | Non-domain Regions |  |  |
| Training Data | $r^2$ | RMSE | $\rho$ | $r^2$ | RMSE | $\rho$ | $r^2$ | RMSE | $\rho$ |
| Archaea | 0.325 | 8.9 | 0.481 | 0.366 | 8.1 | 0.473 | 0.329 | 9.4 | 0.489 |
| Bacteria | 0.909 | 11.9 | 0.829 | 0.926 | 9.4 | 0.816 | 0.893 | 15.0 | 0.833 |
| Both | 0.780 | 5.0 | 0.643 | 0.815 | 4.6 | 0.658 | 0.717 | 5.7 | 0.601 |

Transferability within prokaryotes depends on whether Archaea or Bacteria data is used as the training dataset and is highest when only Pfam domain amino acid composition is used (Figure 2A). However, models that use proteome-wide amino acid frequencies cannot predict temperature sensitivity in eukaryotes, and achieve an average r^2^=0.03 (Figure 2B, Table 2). These models are not, therefore, helpful for understanding eukaryotic protein adaptation to high temperature.

**Figure 2:**
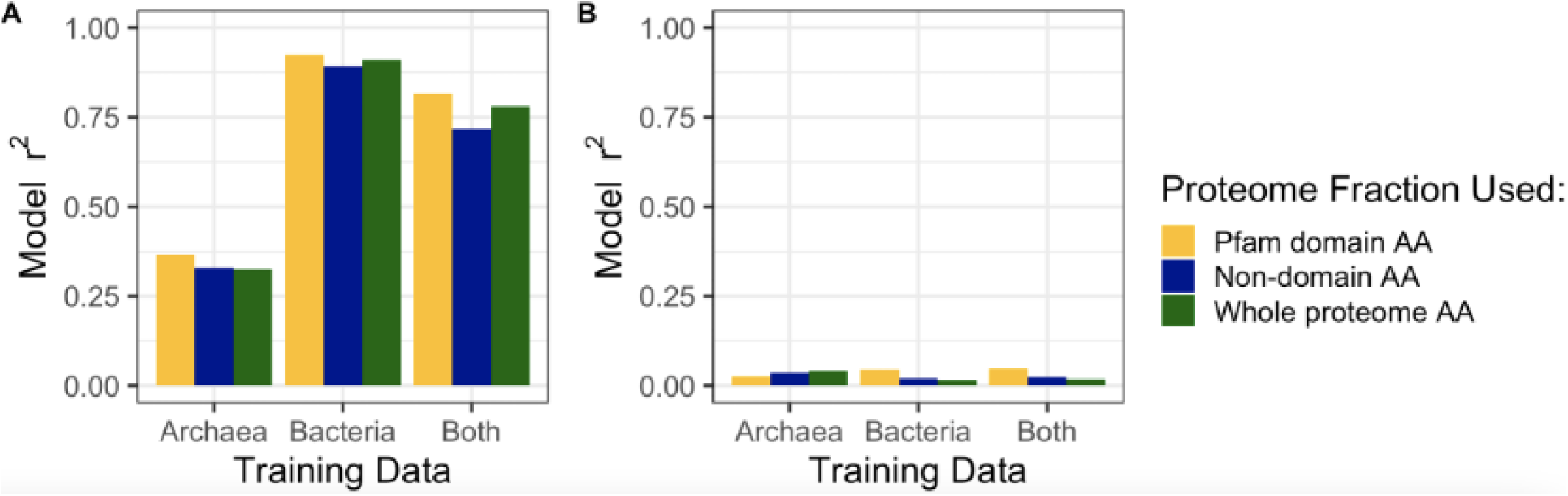
Global amino acid models accurately predict prokaryote optimal growth temperatures, but not eukaryote optimal temperatures. A) Model r^2^ in prokaryotes; B) model performance when applied to six eukaryote species. Colors in both plots reflect the type of input data used: yellow = Pfam domain regions of the proteome, blue = non-Pfam domain regions of the proteome, green = whole proteome. Transferability was tested for six eukaryotic species (Caenorhabditis elegans, OGT=20°C; Arabidopsis thaliana, OGT=25°C; Danio rerio, OGT=28°C; Drosophila melanogaster, OGT=28°C; Saccharomyces cerevisiae, OGT=30°C; Mus musculus, OGT=37°C; eukaryotic OGTs from Jarzab et al. 2020).

**Table 2:** Global model performance when applied to eukaryotic species.

|  | Dataset |  |  |  |  |  |  |  |  |
| --- | --- | --- | --- | --- | --- | --- | --- | --- | --- |
|  | Whole Proteome |  |  | Pfam Domains |  |  | Non-domain Regions |  |  |
| Training Data | $r^2$ | RMSE | $\rho$ | $r^2$ | RMSE | $\rho$ | $r^2$ | RMSE | $\rho$ |
| Archaea | 0.042 | 18.5 | -0.348 | 0.026 | 12.5 | -0.203 | 0.035 | 20.5 | -0.348 |
| Bacteria | 0.016 | 15.8 | -0.348 | 0.046 | 16.6 | -0.348 | 0.019 | 14.3 | -0.348 |
| Both | 0.017 | 15.3 | -0.348 | 0.046 | 16.8 | -0.348 | 0.023 | 15.1 | -0.348 |

### Pfam domains explain temperature sensitivity with variable transferability

Because the most accurate global model uses Pfam domain amino acid frequencies, we asked whether individual Pfam domains can be used to predict protein temperature sensitivity. Pfam domains were identified in each prokaryote proteome and separated into clusters based on k-means clustering of sequence features. 5,283 Pfam domains were identified in prokaryotes and were separated into 2-12 sequence clusters. A ridge regression model was built for each Pfam domain cluster and tested to see how well it could predict prokaryote OGT or eukaryote protein melting temperatures.

There is a range of prediction accuracies within prokaryotes. As with the global models, a joint dataset containing both Archaea and Bacteria data performs best. There were 16,158 unique Pfam domain clusters that could be modeled, resulting in an average model r^2^=0.30 when using the combined Archaea and Bacteria dataset. In this combined dataset, 80% of the data in each Pfam domain was randomly selected and used to build the model and the remaining 20% was held out to evaluate model performance. Fewer Pfam domains could be compared when models were built with only Archaea or only Bacteria species’ data due to the small number of available Archaea species. In both cases, models were built with one phylogenetic domain and accuracy was determined by predicting OGT for the other phylogenetic group (e.g. models were built using Archaea OGTs and Pfam domains were then evaluated based on their ability to predict Bacteria OGTs). In total, 2,619 Pfam domain clusters were tested and compared across these two phylogenetic domains. Models built using Bacteria data had an average r^2^=0.20 when used to predict Archaea OGTs, while models built using Archaea data achieved only r^2^=0.02 when used to predict Bacteria OGTs (Figure 3A; Table 3).

**Figure 3:**
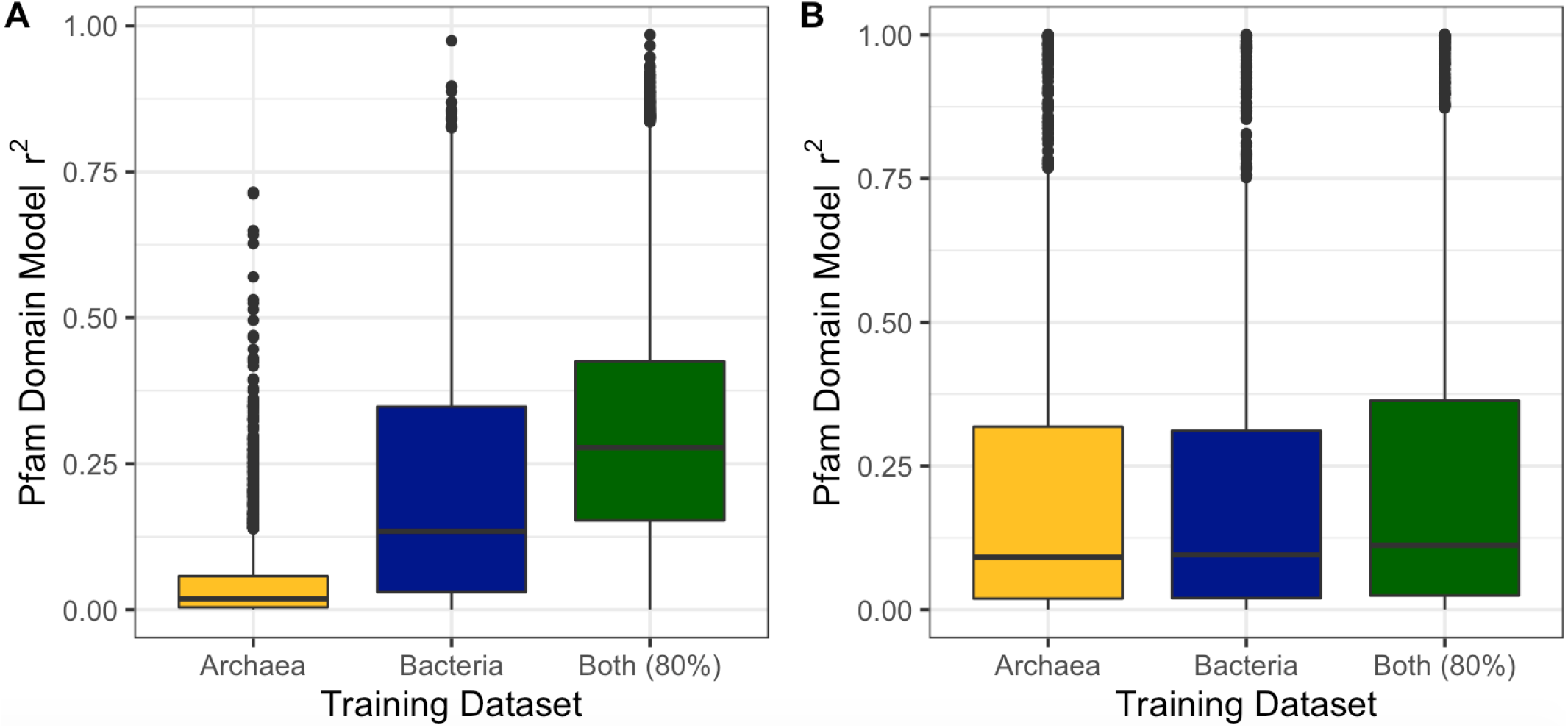
Pfam domain models perform better than global proteome models when applied to eukaryotic species. A) Distribution of model performances across all evaluated Pfam domain clusters when Pfam domain amino acid frequencies are used to predict prokaryote optimal growth temperatures. B) Performance of models when used to predict eukaryotic protein melting temperatures. Colors represent the different model training and test sets: yellow = built with Archaea, tested in Bacteria (or eukaryotes); blue = built with Bacteria, tested in Archaea (or eukaryotes); green = built with 80% of total data, tested with held-out 20% (or eukaryotes).

**Table 3:** Average Pfam domain model performance across training and test sets

| Training Data | Testing Data | Average $r^2$ | Average RMSE | Average $\rho$ |
| --- | --- | --- | --- | --- |
| Archaea | Bacteria | $0.02 \pm 0.044$ | $22.5 \pm 11.2$ | $0.05 \pm 0.15$ |
| Bacteria | Archaea | $0.20 \pm 0.19$ | $21.5 \pm 8.7$ | $0.22 \pm 0.32$ |
| Both (80%) | Both (20%) | $0.30 \pm 0.19$ | $8.18 \pm 4.36$ | $0.20 \pm 0.20$ |
| Archaea | Eukaryote | $0.34 \pm 0.38$ | $16.0 \pm 10.8$ | $0.02 \pm 0.41$ |
| Bacteria | Eukaryote | $0.34 \pm 0.38$ | $15.3 \pm 6.1$ | $0.04 \pm 0.42$ |
| Both (80%) | Eukaryote | $0.38 \pm 0.39$ | $16.1 \pm 5.7$ | $0.02 \pm 0.42$ |

The same models were then applied to eukaryotic species to see whether they could effectively transfer information across the prokaryote-eukaryote divide. These Pfam domain models transfer to eukaryotic species better than global amino acid frequency models and are able to predict eukaryotic protein melting temperatures (T_m_) with average r^2^ = 0.34-0.38 (Figure 3B). However, none of the correlations are significant at an FDR significance threshold of 0.05 and Spearman’s ρ values are low when Pfam domain models were applied to eukaryotic Pfam domains, suggesting that Pfam domain models still cannot reliably transfer stability estimates across the prokaryote-eukaryote divide (Table 3).

Pfam domain models range in accuracy, with amino acid composition being highly predictive of OGT for some domain models but not for others. Only 450 Pfam domains can be associated with Gene Ontology (GO) terms. GO enrichment analysis shows that the most transferable models - models that rank eukaryotic proteins with r^2^ > 0.5 - were slightly enriched for GO terms related to metabolism and catalytic activity (Supplemental Figures 1-2).

### Over one-third of the proteome is associated with species’ optimal temperature

The large number of prokaryotic sequences available for most Pfam domains makes it possible to associate individual sites with prokaryote optimal growth temperatures. For each Pfam domain cluster, domain observations were aligned and re-coded to reflect each of 9 amino acid physicochemical properties (Li et al. 2016). Two principal components were added to control for species relatedness; these explain an average of 94% of the variance within each domain cluster (Supplemental Figure 3). Site-specific associations were run separately in Archaea and Bacteria species, and the resulting positions were compared to determine whether sites could be consistently associated with OGT across phylogenetic domains.

323,850 residue positions in 2,088 Pfam domain clusters were tested for associations with species’ OGT. Associations were calculated separately in Archaea and Bacteria species and OGT-associated sites were compared between the two phylogenetic domains to determine whether a consistent subset of residue positions are associated with optimal temperature. On average, 59.0% of sites in Archaea Pfam domains and 75.2% of sites in Bacteria Pfam domains pass an FDR significance threshold of α=0.05. There is a substantial overlap between sites in Archaea and Bacteria, with 48.0% of Pfam domain positions associated with temperature in both phylogenetic domains (Figure 4A). The lower proportion of significant sites in Archaea Pfam domains may be due to reduced power to detect associations in the smaller Archaea dataset. There is a positive rank correlation between p-values from sites in the Archaea dataset and sites in the Bacteria dataset for most Pfam domains, and the correlation is significant in nearly 40% of the tested Pfam domain groups, suggesting that similar positions within a protein domain are temperature sensitive in both Archaea species and Bacteria species (Figure 4B). For most of the significant sites, 4-7 physicochemical properties have consistent effect estimates and this distribution differs significantly from the expected distribution of effect estimates (Χ-square test, p < 2.2e-16). The set of physicochemical properties that has a consistent effect estimate in both Bacteria and Archaea GWAS results varies from site to site, but each property is consistent across domains in about 50% of the tested positions (Supplemental Figure 4).

**Figure 4:**
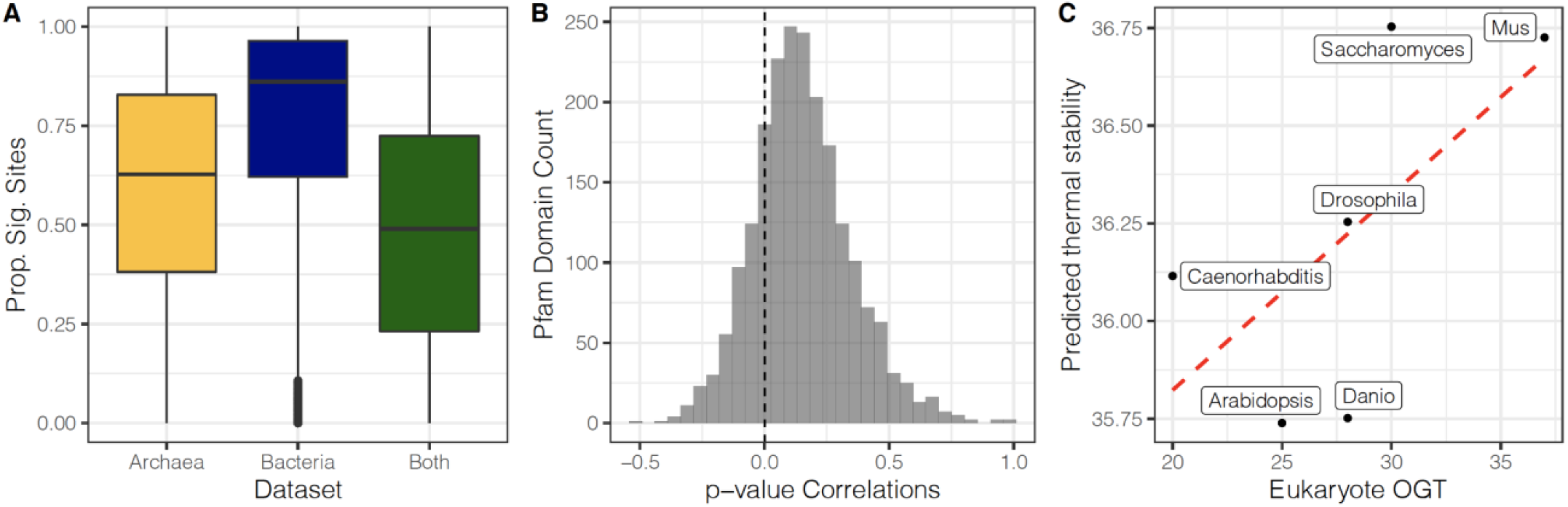
Individual Pfam domain sites are associated with temperature. A) Proportion of Pfam domain sites that pass a 5% FDR significance threshold in either Archaea (yellow), or Bacteria (blue), or are shared between both (green). B) Correlation of p-values between Archaea associations and Bacteria associations for each Pfam domain cluster. C) Relationship between the true optimal growth temperature of six eukaryote species and the predicted thermal stability in these species. Thermal stability is predicted using residue-specific associations with optimal temperature for single-copy orthologous proteins present in all six species (outlined in Figure 4.8).

**Figure 5:**
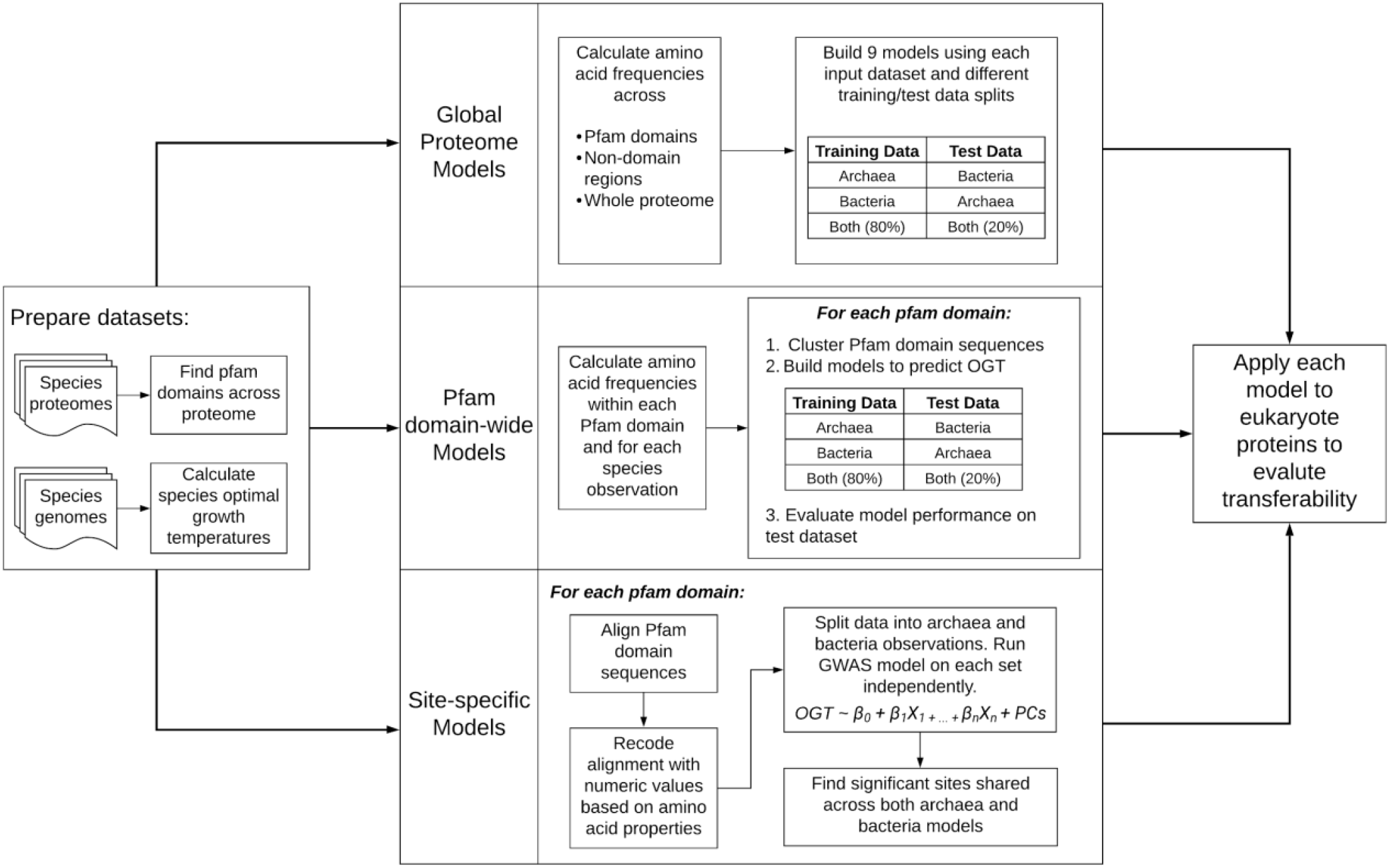
General steps for each model to evaluate proteome composition as it relates to temperature. Briefly, species’ proteomes and genomes are processed to predict OGT and to find Pfam domains. This data is used to create models predicting temperature sensitivity using global proteome features, Pfam domain features, or site features.

**Figure 6:**
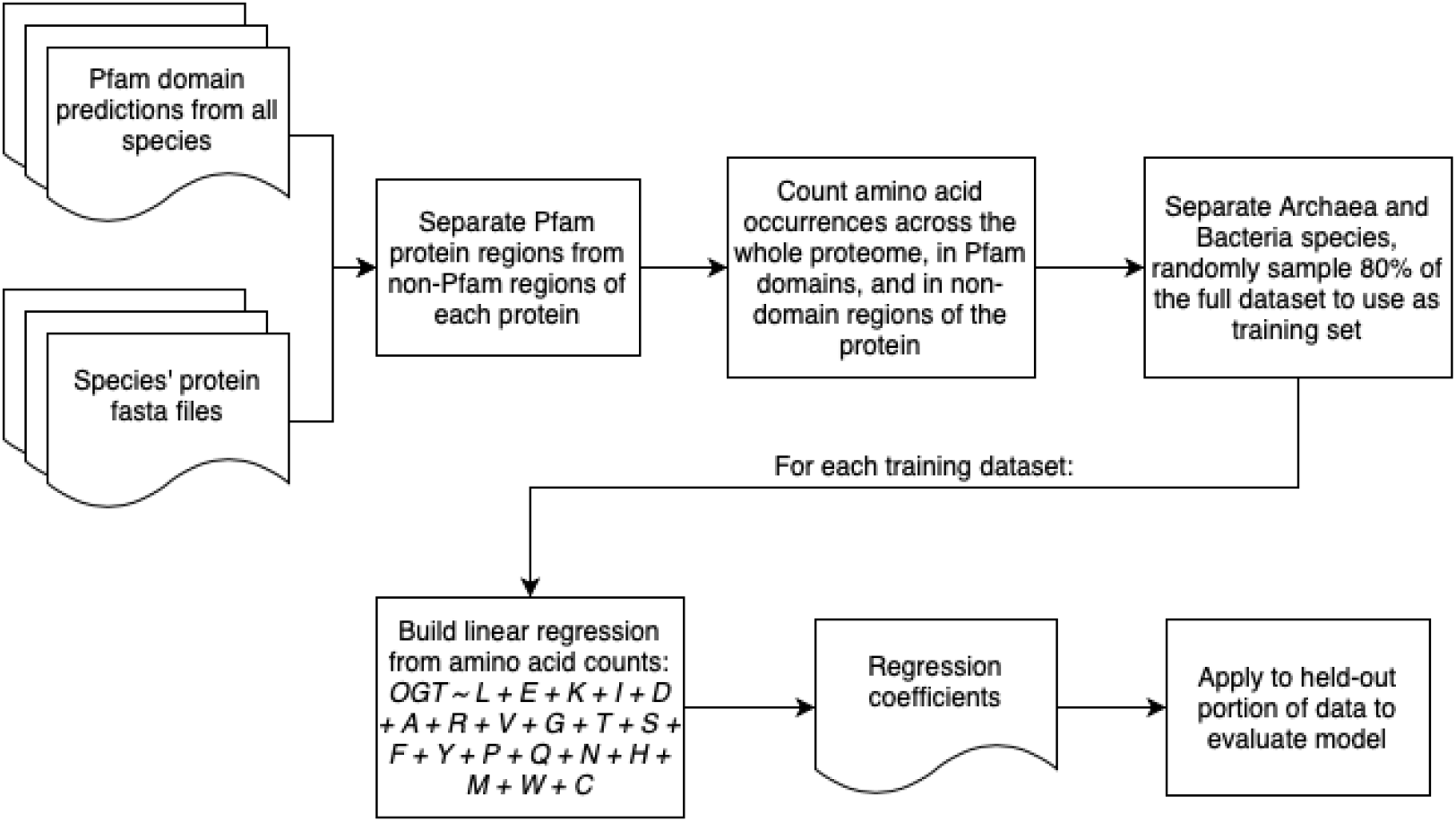
Logic used to build global proteome models to predict OGT from species’ amino acid counts.

**Figure 7:**
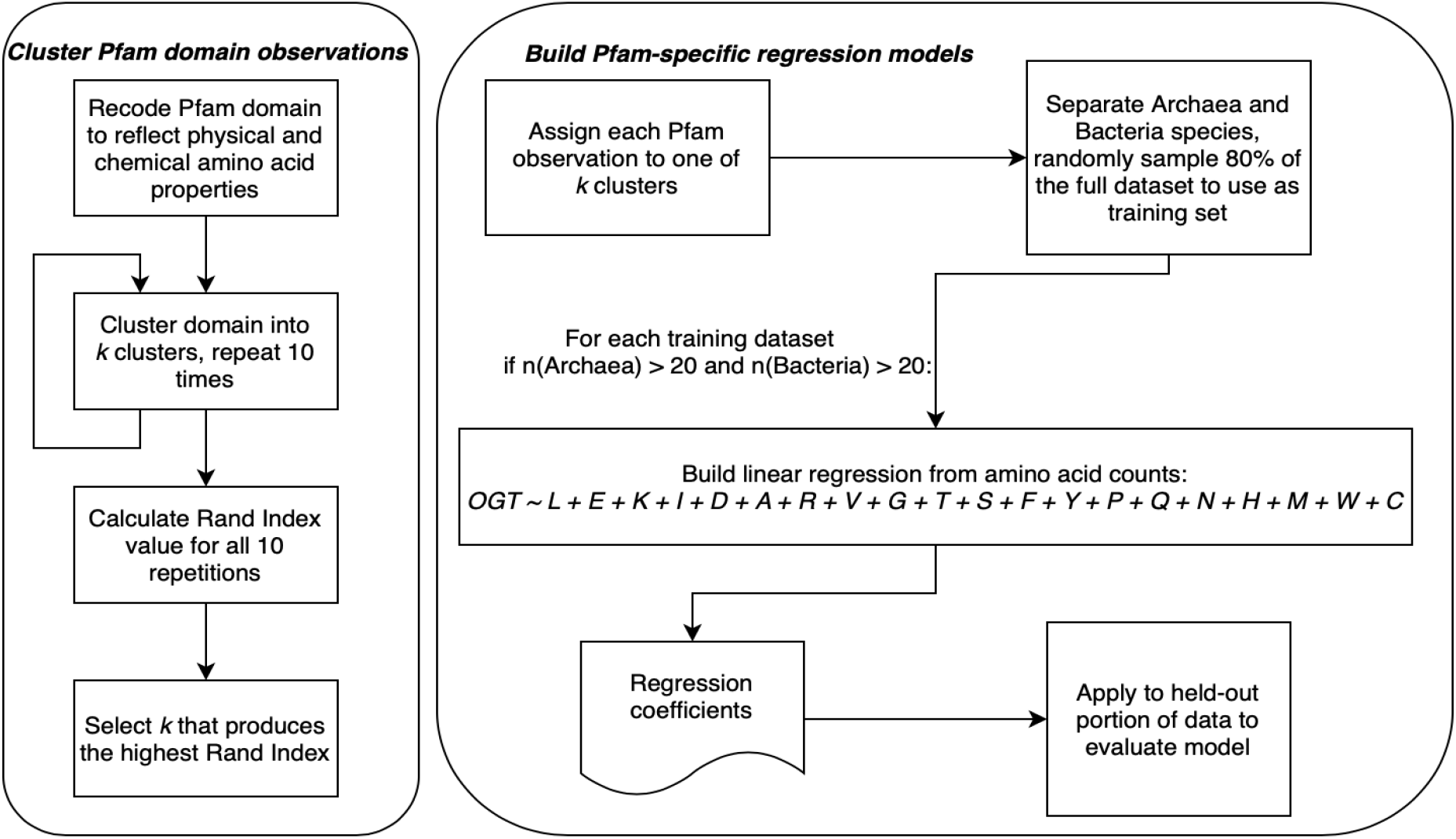
Logic used to cluster Pfam domains and build domain-specific regression models to predict optimal temperature.

**Figure 8:**
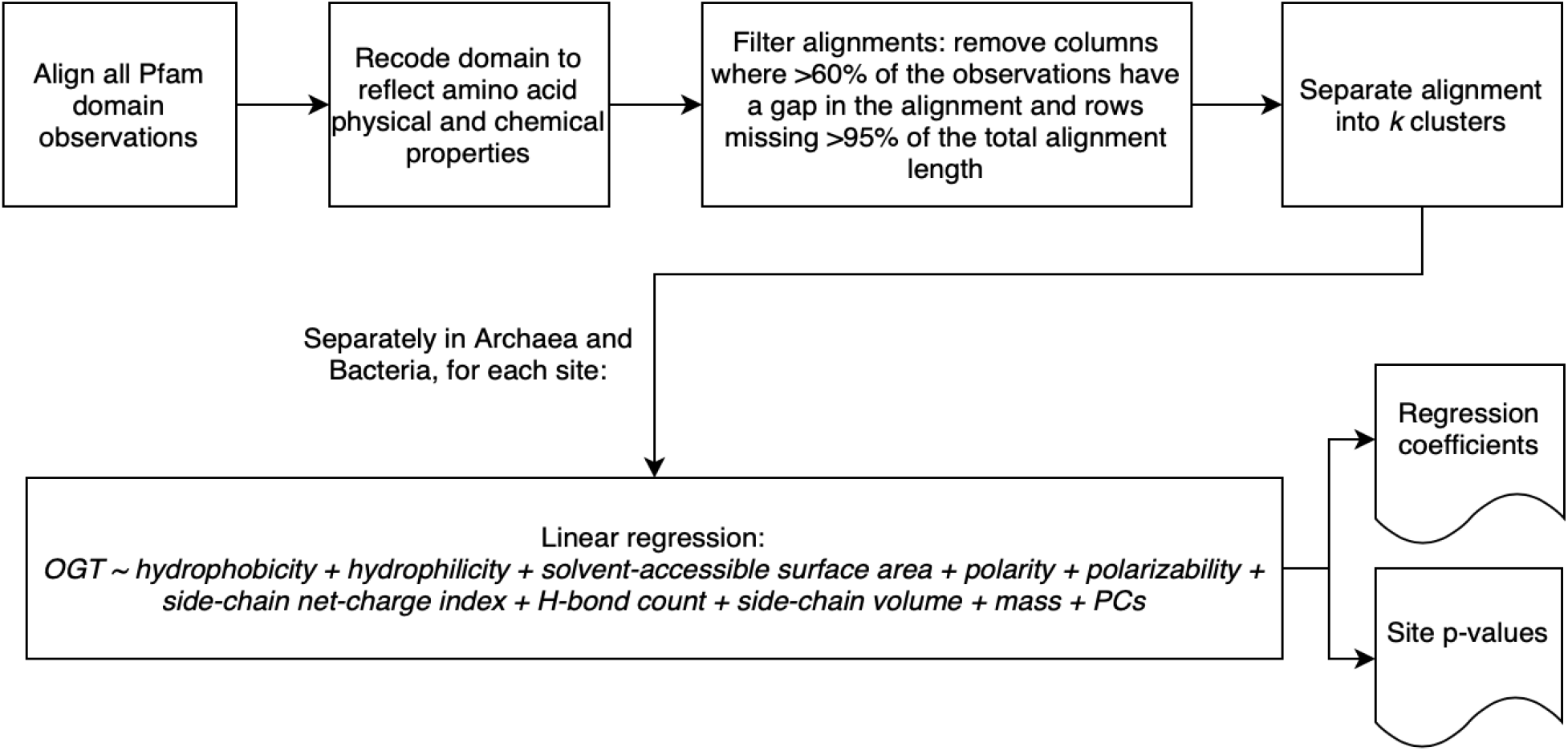
Logic used for site-specific GWAS models on Archaea and Bacteria Pfam domains.

To compare the transferability of this site-specific model to eukaryotes, we identified 41 single-copy orthogroups present in six eukaryotes. We used the prokaryote dataset to identify temperature-associated residues and calculated a value of temperature sensitivity for each site based on the mean OGT of prokaryotic species with the same amino acid at the same position. Temperature sensitivity values were then aggregated across all tested proteins. The correlation between the resulting measure of thermal sensitivity and the true OGT for the six eukaryotic species is r^2^=0.39 (p=0.18) and the rank correlation is ρ= 0.72 (p=0.10). This is suggestive of an improvement over both the global proteome and Pfam domain models, although with so few eukaryotic species the correlation is not significant (Figure 4C).

## Discussion

In this study, we evaluated protein sequences from more than 4,800 prokaryotic species to investigate whether trends in protein sequence or composition could be associated with optimal growth temperature and applied across a wide range of species. Our aim was to build transferable models of temperature adaptation for species from any phylogenetic domain using primary amino acid sequence. Our analyses show that while proteome-wide counts are good predictors of prokaryotic OGT, extension to eukaryotic species requires comparisons using smaller sections of the proteome. Protein domains, which are highly conserved across the tree of life, are a good candidate for building transferable models across large phylogenetic distances to understand trends in molecular evolution, and GWAS-like models can identify individual residues that are associated with optimal growth temperature.

While we were able to replicate findings from previous studies correlating amino acid frequency in prokaryotes with OGT (Zeldovich et al. 2007; Sauer and Wang 2019), these models cannot predict OGT for eukaryotes. We hypothesize that the failure to transfer global proteome models to multicellular eukaryotes is due to the additional complexity, lower effective population sizes, and longer generation times of the eukaryotic species tested here. These forces reduce the efficiency of selection on individual amino acid residues within eukaryotes (Huber et al. 2017). As a result, amino acid frequencies across the proteome are more influenced by genetic drift and do not accurately reflect temperature optima in eukaryotes the way they do in prokaryotes.

For global models that incorporate information from across the proteome, Pfam domain regions of the protein are more predictive than the proteome as a whole. Models using Pfam domain amino acid frequencies to predict OGT can also predict eukaryote Tm with moderate levels of accuracy. Previous studies have found that protein functional requirements constrain evolution and amino acid frequency, which is consistent with our findings that functional Pfam domain regions are more predictive of prokaryote OGT than the proteome as a whole (Saelensminde et al. 2007; Zeldovich et al. 2007; Arcus et al. 2016). Since Pfam domains often carry out important binding and catalytic functions, it seems likely that these sequences are under strong selective pressure and that deleterious temperature-sensitive mutations within Pfam domains would be purged quickly (El-Gebali et al. 2019). Selection may be weaker in the loop and disordered protein regions outside of Pfam domains, increasing noise and reducing the signal of temperature sensitivity.

The site-specific models that associate individual amino acid residues with temperature give insight into which positions within a protein are temperature sensitive. This model also transfers best across the prokaryote-eukaryote divide, although it can only rank species by temperature sensitivity and cannot predict optimal temperature directly. These models likely transfer to eukaryotes better than models based on amino acid frequencies because they consider each residue separately, and only use residues associated with OGT in prokaryotes to make predictions about eukaryotic amino acid residues. The results support our hypothesis that a consistent subset of amino acid residues is important for protein evolution, and that many small changes in aggregate contribute to protein evolution and thermal adaptation (Petrović et al. 2018; Venev and Zeldovich 2018).

This method provides the community with a rich set of predictions about which amino acids are temperature sensitive in Pfam domains and establishes methods to evaluate amino acid temperature sensitivity in other species. Estimates of temperature sensitivity can be used to provide context for amino acid changes across a range of species. For example, a single amino acid change in the maize HPC1 gene has been related to differences in phospholipid composition in warm- and cold-adapted maize varieties. That amino acid change is present in the phospholipase Pfam domain (PF01764) and is significantly associated with OGT in prokaryotes, suggesting that this residue has a consistent and important effect on protein function at different temperatures across both prokaryotes and eukaryotes (Rodríguez-Zapata et al. 2021).

Further research is still needed to relate protein temperature sensitivity to organism fitness and to be able to predict how much a specific amino acid change will increase or decrease thermostability in a eukaryotic protein. Future models may also incorporate linkage or residue interactions to understand how a protein evolves (Petrović et al. 2018; Salinas and Ranganathan 2018). In this study we restricted most of our analyses to Pfam domains to make it easier to identify and align sequences across large evolutionary distances, but loop and disordered protein regions may play an important role in protein temperature sensitivity and warrant further study.

Despite these limitations, the results presented here demonstrate that some biochemical features are transferable across all three domains of life. We show that models focused on small regions of the proteome outperform global models based on amino acid frequency, and that protein evolution in prokaryotes can be used to gain insight into eukaryotic proteome thermal profiles. The methods developed here can be applied to newly-sequenced eukaryotic proteomes and will facilitate research into how protein temperature sensitivity interacts with organism fitness, allow comparisons of molecular temperature sensitivity across species, and help prioritize functional variants when determining candidate mutations for genome editing.

## Materials and Methods

### Selecting species and determining OGT

All prokaryote species with both genomes and proteomes available on EnsemblBacteria were downloaded for evaluation, resulting in a total of 44,048 genomes with some species having many duplicate genomes. We used tRNAscanSE to identify tRNAs in each genome. When there were multiple genomes available, only the genome with the highest number of tRNAs identified was selected. This resulted in a set of 4,827 unique species with both genome and proteome data, including 277 Archaea species and 4,550 Bacteria species. Optimal growth temperature (OGT) was predicted for each species, using the model outlined in Cimen, Jensen, and Buckler (2020). Eukaryotic species with protein melting temperatures were obtained from the meltome atlas dataset (Jarzab et al. 2020).

### Identifying Pfam domains

The PfamScan pipeline was used to predict Pfam domains within each species proteome, and results were then sorted and separated into distinct files for each Pfam domain in the Pfam database (El-Gebali et al. 2019). Pfam domains were aligned using the hmmalign tool from HMMER and default parameters (hmmer.org; version 3.3.1), and alignments were re-coded to reflect amino acid physicochemical properties (Li et al. 2016). A Snakemake pipeline facilitated replication and made it easier to scale the pipeline to many proteomes (Köster and Rahmann 2012). The number of observations varied by multiple orders of magnitude, with some Pfam domains having more than 100,000 observations, and others having no observations at all. To repeat this step, a docker image is available on Dockerhub at lynnjo/protein_temp:0.07 and scripts are available in a Bitbucket repository at https://bitbucket.org/bucklerlab/p_proteintemp/src/master/.

### Global proteome models

The global OGT prediction models require amino acid frequency statistics from across each species’ proteome. Sites were grouped by whether or not they fell within Pfam domains. Amino acid frequencies were calculated for each proteome. Pfam domain amino acid counts were subtracted from the total proteome amino acid counts to determine amino acid frequency in non-Pfam domain protein regions. To see if previous literature results could be replicated, we used a linear regression to predict OGT from amino acid frequency statistics (Figure 1, Table 1). Models were built using data from only Bacteria species, only Archaea species, or a randomly-selected 80% of the total dataset, including both Archaea and Bacteria. In each case, the model transferability was tested by applying the regression model coefficients to the held-out portion of the dataset (Archaea, Bacteria, and 20% of total, respectively). Performance was evaluated by comparing Spearman rank correlation (ρ) and r^2^ between models. Models with higher average performance on the held-out set were considered more transferable. Regression coefficients were then applied to amino acid counts from six eukaryotic species to predict eukaryote OGT and evaluate performance across the prokaryote-eukaryote divide.

### Pfam domain models

#### Clustering domain observations

Pfam domains represent conserved functional segments of proteins, but similar proteins with slight functional differences are often grouped within a single Pfam domain (e.g. PF00006 contains both alpha and beta subunits of ATP synthase). To capture these differences, each re-coded Pfam domain was clustered using the k-means clustering algorithm so that each Pfam domain contained 2 or more clusters. Because kmeans clustering can only be done on numerical values, each Pfam domain amino acid sequence was re-coded to reflect chemical properties of each amino acid before clustering (Li et al. 2016). Each domain was tested with 2 < *k* < 12 clusters. For each value of *k*, the clustering was repeated 10 times with random starting seed values. The Rand Index, which measures similarity between a set of data clusters, was used to evaluate clustering consistency at each value of *k*. The smallest value of *k* that produced reliable data clustering was selected (Rand 1971). Each observation in every Pfam domain was assigned a cluster ID that was used for subsequent analyses.

#### Pfam domain regression analysis

Pfam domain clusters with fewer than 20 Archaea and 20 Bacteria observations were removed. The remaining domains were used to build ridge regression models for each Pfam domain. Models attempted to predict OGT from amino acid composition within the Pfam domain and were then applied to a held-out portion of the data to determine how well information could be transferred across domains of life. As with the global models, data were either split by phylogenetic domain (e.g. trained in Bacteria, applied to Archaea and vice-versa) or by randomly selecting 80% of the data to build the model and predicting OGT for the remaining 20%. Predicted OGT was compared to known OGT to determine model accuracy. Prediction r^2^ and root mean square error (RMSE) were used to evaluate performance.

After building and comparing model performance on prokaryotic Pfam domains, the same models were applied to eukaryotic proteins from the meltome atlas dataset (Jarzab et al. 2020). Pfam domains were predicted for six eukaryotic species from the meltome atlas using the Pfam prediction Snakemake pipeline, and amino acid frequencies were calculated for each domain as with prokaryotes. Prokaryotic protein domain model coefficients were used to predict T_m_ for eukaryotic protein domains and compared to the measured T_m_ values from the meltome atlas.

#### GO analyses

GO terms associated with transferable Pfam domain models were evaluated using the topGO program in R (Alexa and Rahnenfuhrer 2020). Models were considered transferable if they resulted in an r^2^ > 0.5 when applied to eukaryotic protein melting temperatures.

### Site-specific models

Pfam domain observations were aligned using hmmalign (hmmer.org; version 3.3.1) using default parameters. Alignments were clustered using the method described above, and re-coded to reflect amino acid physicochemical properties (Li et al. 2016). Each Pfam domain was filtered to remove columns in the alignment that contained a gap character at 60% or more of the observations. The remaining alignment columns were then filtered to remove rows that were missing more than 95% of the total Pfam domain sequence, as these Pfam domain fragments are unlikely to be functional domains (Triant and Pearson 2015).

There were 1,243 Pfam domains with an average of 260.5 observations per domain left after filtering, resulting in a total of 323,850 tested residue positions. Pfam domain lengths vary from species to species, but the same set of 1,243 Pfam domains contained 801,168 positions in the aligned Pfam sequences, meaning that 40% of sites were retained after filtering. For each cluster and position in the Pfam domain, a separate linear regression was run to associate the residues at that position with species’ OGT. Terms included numerical values representing amino acid hydrophobicity, hydrophilicity, hydrogen bond count, side-chain volume, polarity, polarizability, solvent-accessible surface area, side chain net charge index, and mass, as well as 2 principal components (PCs) to account for phylogenetic relationships between species within the cluster. PCs were calculated on the filtered sequence data and used to control for shared evolutionary history between species. Initial tests of variance explained by 1-3 principal mcomponents showed that most of the variance in amino acids at a particular site could be captured by a single principal component, with an average of 92% of the variance within the domain sequences explained by one principal component and 94% and 95% explained by 2 and 3 PCs, respectively (Supplemental Figure S3).

To determine whether similar residues were responding to temperature in both Archaea and Bacteria, site-specific regressions were run in each phylogenetic domain separately. Sites that passed a 5% Bonferroni-corrected significance threshold in both Archaea and Bacteria were compared. The Spearman rank correlation of p-values in the Archaea and Bacteria analyses were used to determine whether OGT associations were similar across the two phylogenetic domains.

Six eukaryotic species were used to determine whether sites associated with temperature in prokaryotic species were also predictive of eukaryotic temperature adaptation. Single-copy orthogroups were identified with OrthoFinder, resulting in 41 single-copy orthogroups shared among all six species (Emms and Kelly 2015). Pfam domains were identified in the shared orthogroup proteins and aligned to the existing prokaryote Pfam domain sequences using the mafft --add function and default parameters, which ensured that alignment coordinates with the added eukaryotic sequences matched coordinates from the original prokaryote alignments (Katoh and Frith 2012). At each site, the eukaryotic amino acid residue was compared to the average OGT of prokaryote species with the same residue at that position, giving a measure of thermal adaptation for each site in every Pfam domain.

Because the Pfam domains identified in homologous proteins can vary across species, a different number of sites was tested for each of the six eukaryotes (Table 4). All sites that were significantly associated with OGT in prokaryotes (and passing an FDR significance threshold of 0.1) were then collected and averaged across all sites to get a statistic representing thermal adaptation for each species. Sites were averaged to provide a single number for each genome for comparison purposes. The distribution of predicted OGT values was similar in all six species, with bimodal normal distributions centered around 30°C and 50 °C. Both r^2^ and ρ decrease if using mode OGT values instead of averages to account for the bimodality; r^2^=0.17 and ρ=0.64. This thermal adaptation statistic was compared to the overall OGT of the species to evaluate transferability.

**Table 4:** Testable amino acid residues and Pfam domain count for six eukaryote species.

| Species | Pfam domain count | Residue count | Average sites/domain |
| --- | --- | --- | --- |
| *Arabidopsis thaliana* | 34 | 5912 | 173.9 |
| *Caenorhabditis elegans* | 62 | 9000 | 145.2 |
| *Danio rerio* | 31 | 4836 | 156.0 |
| *Drosophila melanogaster* | 37 | 5745 | 155.3 |
| *Mus musculus* | 30 | 5084 | 169.5 |
| *Saccharomyces cerevisiae* | 28 | 5000 | 178.6 |

## Data availability

The results presented in this paper can be found on the CyVerse Data Commons at /iplant/home/shared/commons_repo/curated/Jensen_proteinTemp_Jun2021. Scripts and the snakemake pipeline developed to facilitate the analyses can be found on bitbucket at https://bitbucket.org/bucklerlab/p_proteintemp/src/master/.

