## Supplemental Figures 1-4 for "Predicting protein domain temperature adaptation across the prokaryote-eukaryote divide"

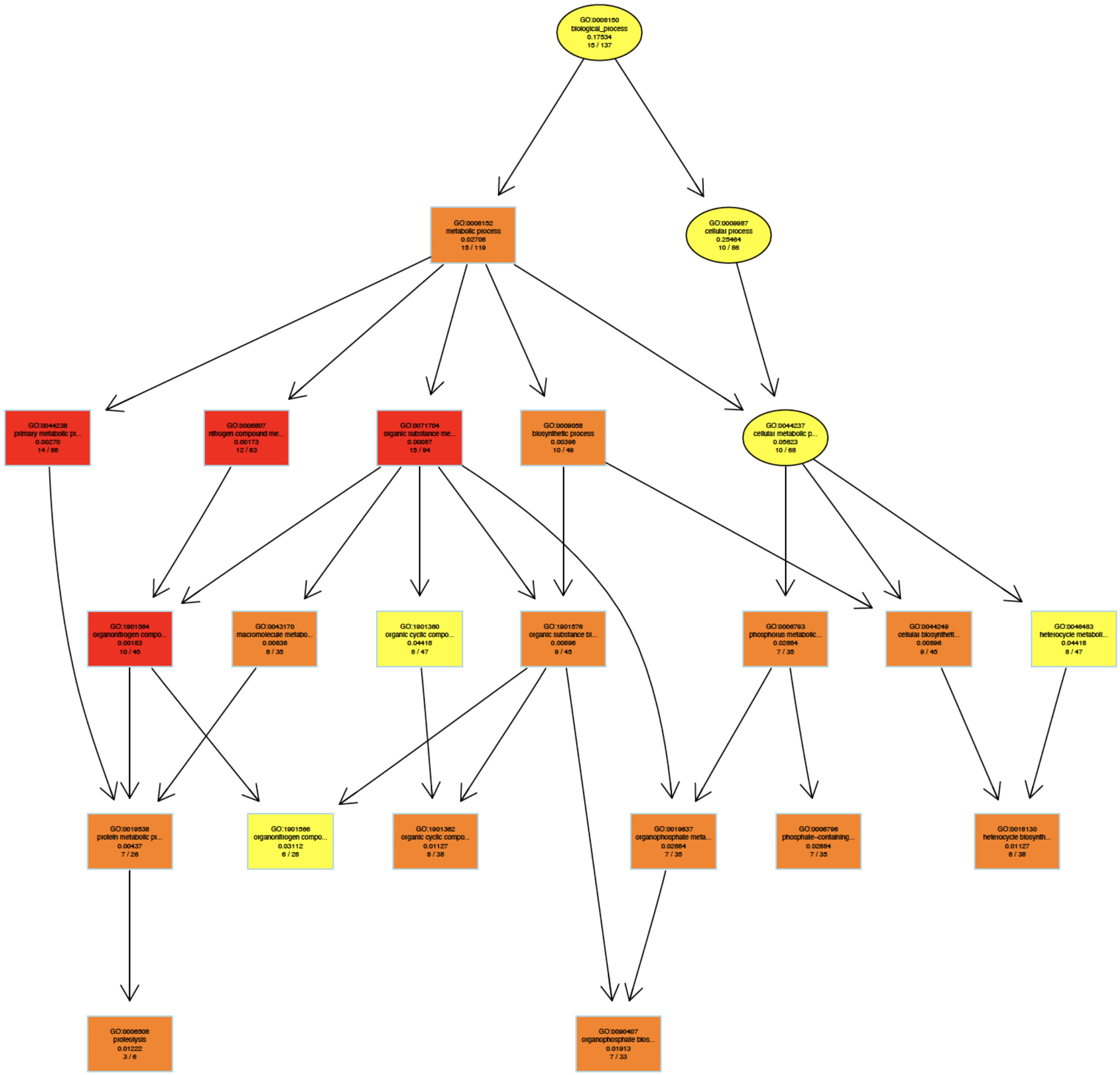


**Supplemental Figure 1:** GO terms enrichment showing the top Biological Process terms identified by the classic algorithm for enrichment. Boxes include the terms that are significant at the level of $p<0.05$, and box color shows the relative significance with dark red indicating the most significant terms and light yellow indicating the least significant terms. Arrows indicate is-a relationships between GO terms.

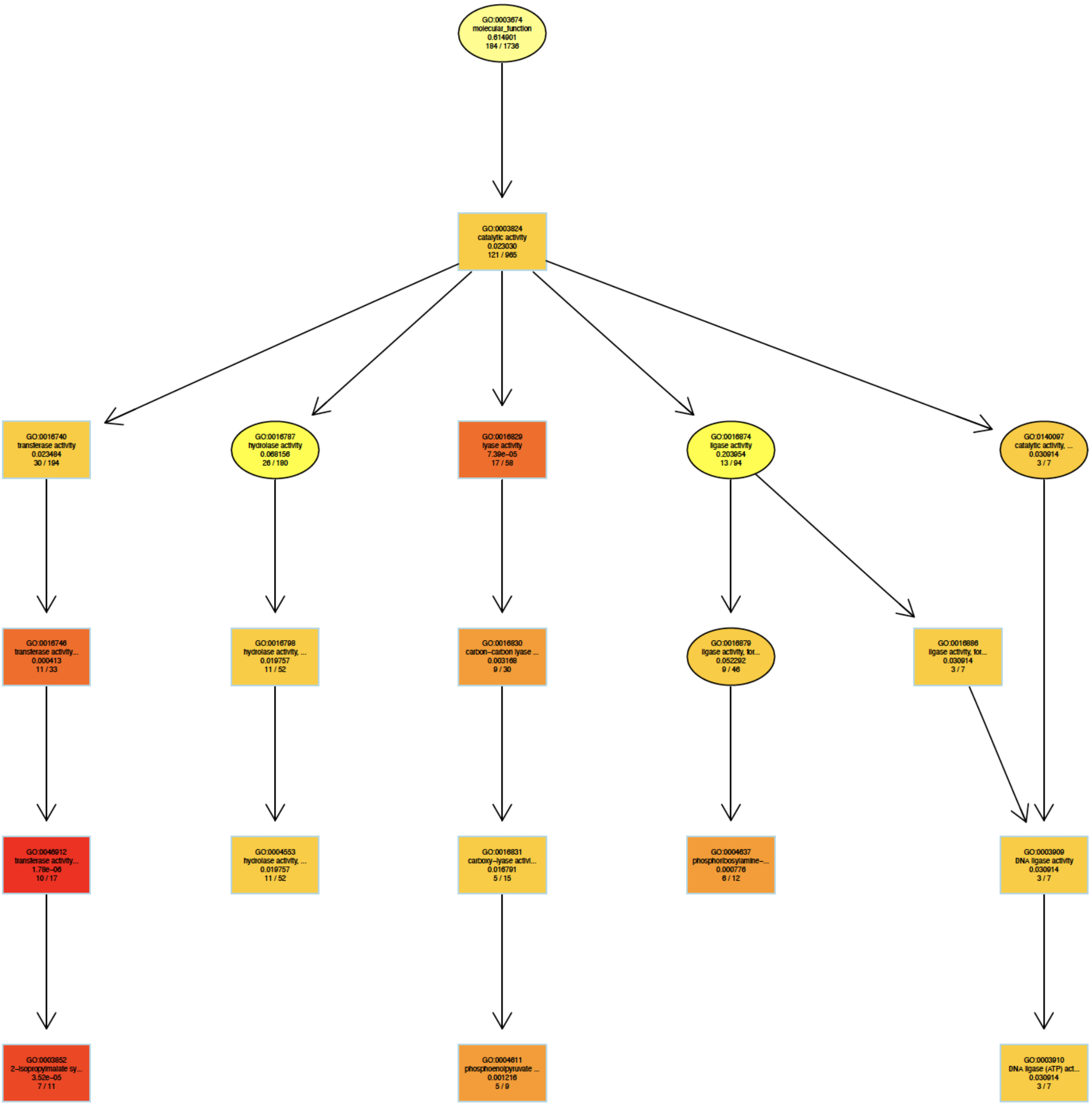


**Supplemental Figure 2:** GO terms enrichment showing the top Molecular Function terms identified by the classic algorithm for enrichment. Boxes include the terms that are significant at the level of $p<0.05$, and box color shows the relative significance with dark red indicating the most significant terms and light yellow indicating the least significant terms. Arrows indicate is-a relationships between GO terms.

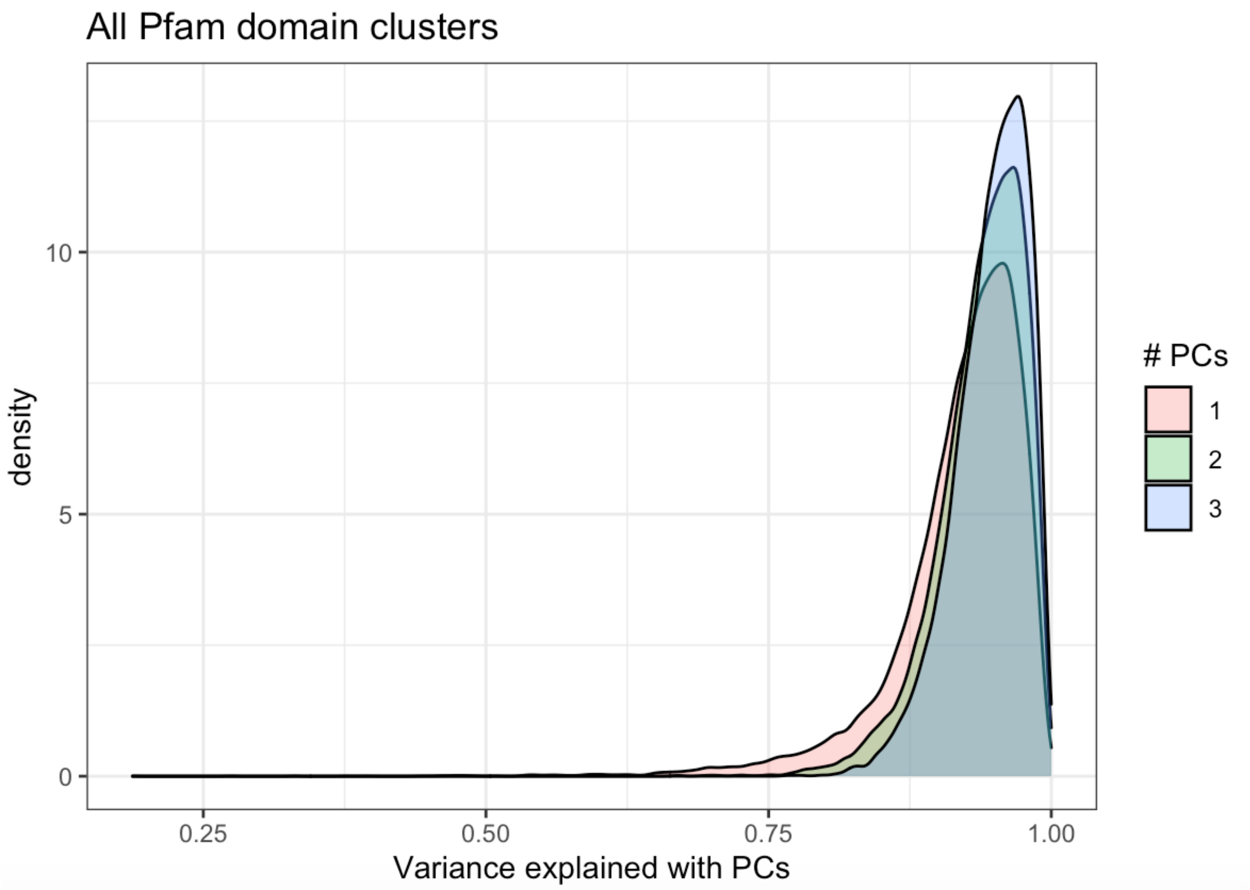


**Supplemental Figure 3:** Proportion of variance within each Pfam domain captured by different number of principal components. Pink = one principal component, green = two principal components, blue = three principal components.

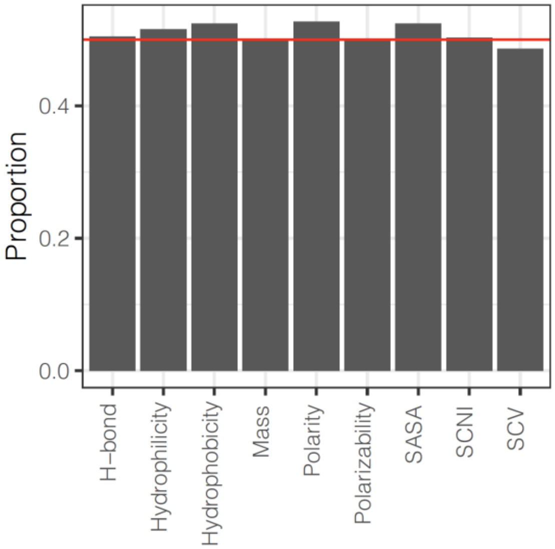


**Supplemental Figure 4:** Effect estimates of each physicochemical property effect estimates have a consistent sign in about 50% of both Archaea and Bacteria GWAS hits. For comparison purposes, a red line is included at y=0.5.
